# *CROCKETA:* An automated framework for comprehensive multi-omic analysis of gene expression and clonotype immune repertoire at a single-cell level

**DOI:** 10.1101/2025.05.16.654451

**Authors:** Gonzalo Soria-Alcaide, Gonzalo García-Aguilera, Marta Portasany-Rodriguez, Jaanam Lalchandani, África González-Murillo, Elena G. Sánchez, Cristina Saiz-Ladera, Manuel Ramírez-Orellana, Jorge Garcia-Martinez

## Abstract

*CROCKETA* (single-**C**ell **R**epertoire **O**rganization & **C**ombined **K**inetics **E**xploration for **T**ranscriptomic **A**nalysis) is an automated and adaptable *Snakemake* pipeline designed to perform fundamental initial stages of single-cell analysis for both transcriptomic and immune repertoire data, with additional steps for detailed assay characterization. *CROCKETA* stems from the imperative need to devise an optimal methodology for integrating and analyzing single-cell RNA sequencing (scRNA-seq) alongside single-cell T-cell Receptor (scTCR-seq) or B-Cell Receptor (scBCR-seq) data in an automated way by means of state-of-the-art tools. This pipeline encompasses basic steps from primary & secondary scRNA-seq analysis (Quality Control -QC-, sequence alignment, cell-level QC and doublets removal, data preprocessing, cell clustering, cell annotation at both single cell and cluster levels, differential expression analysis, trajectory inference and functional enrichment) along with the incorporation and analysis of immune repertoire data for both B-cells and T-cells, capable of accommodating both human and mouse datasets. The analysis can be conducted from either FastQ-formatted raw data or from expression matrices itself. *CROCKETA* opens up new horizons of possibilities by providing a reproducible, automated, and efficient solution for processing large volumes of data, addressing a challenge that had yet to be resolved in single-cell analyses.

## INTRODUCTION

Lymphocytes are central players in the adaptive immune system, and while they have been widely explored from a transcriptional perspective, critical areas such as lymphocyte clonality remain overlooked. The immune repertoire is comprised of a vast diversity of B-cell receptors (BCRs) and T-cell receptors (TCRs). BCRs and TCRs play a central role in the immune system’s recognition of pathogens as well as antigens, and their analysis can provide valuable insights into areas such as treatment efficacy in cancer and other diseases [1–3]. An immense clonal diversity arises from *V(D)J* recombination, which results in the hypervariable antigen-binding complementarity-determining region 3 (CDR3). Diversity is further amplified by certain genetic mechanisms such as imprecise gene segment junctions or somatic hypermutation in the case of B-cells. The outcome is a potential diversity of over 10^20^ unique TCR immune receptor sequences and 10^18^ unique BCR sequences [4,5]. The collection of TCRs & BCRs within a specific individual constitutes its immune repertoire. The immune repertoire reflects both the immune history and current status of the individual [6–9]. Any threat will be addressed by clonal expansion and selection of B-cells and T-cells [10]. The repertoire is highly dynamic and adaptive, and its study sheds light on the disease prognosis and therapeutic outcomes [11]. CDR3 constitutes the identifier of individual lymphocyte clonality: lymphocytes with identical CDR3 regions are defined as belonging to the same clone [12].

Recent years have seen rapid and unprecedented advancements in scRNA-seq technologies, both in technical capabilities and analytical methodologies. Continuous improvements in sensitivity, resolution and scalability have revolutionized transcriptomics, enabling a more precise exploration of cellular heterogeneity [13]. Concurrently, high throughput sequencing technologies have enabled the exponential development of methods for immune repertoire accession and analysis, leading to a significant increase in the demand for optimal approaches to large-scale clonal analysis [14–16]. These advancements now provide the opportunity to integrate both data sources, allowing for single-cell level insights and exploration of new research horizons. The integration of these two technologies facilitates the investigation of immune transcriptional heterogeneity throughout the disease process while accounting for clonality [17].

However, the absence of gold standards in the analytical domain continues to restrict the reproducibility of transcriptional and clonal analyses. In recent years, scRNA-seq analysis has faced substantial challenges due to the lack of benchmarking frameworks and the methodological heterogeneity in the field [18]. Current approaches to immune repertoire analysis often rely on fragmented pipelines, limiting seamless integration between transcriptional profiling and clonal characterization. While tools such as *Bollito* [19], *platypus* [20], and *screpertoire* [21] offer partial solutions, none provide a unified workflow that enables efficient integration of single-cell data while maintaining the flexibility to accommodate user-specific analytical strategies. Moreover, most existing tools lack state-of-the-art methodologies for the combined analysis of transcriptional and clonal dynamics.

Here, we introduce *CROCKETA* (single-Cell Repertoire Organization & Combined Kinetics Exploration for Transcriptomic Analysis), the first tool to fully integrate transcriptional and clonal analysis within a *Snakemake*-based workflow, enabling large-scale parallelization, exact reproducibility, and modular customization. *CROCKETA* facilitates the exploration of clonal expansion, diversity, and overlap in the context of gene expression within the *Seurat* [22] framework. Additionally, it incorporates cutting-edge methodologies such as *Immunarch* [23] for viral clone annotation, *SingleR* [24] for cell annotation, *Vision* [25] for transcriptional signature analysis or *Slingshot* [26] for trajectory inference, combining capabilities that have not previously been available within a single tool.

Crucially, *CROCKETA’s* ability to correlate transcriptional states with clonal expansion has direct implications for cancer immunotherapy, where identifying expanded clones associated with exhaustion or memory phenotypes could inform the development of cellular therapies. By providing a fully automated, scalable, and integrative solution, *CROCKETA* represents a significant advance in single-cell immune repertoire analysis.

## MATERIALS & METHODS

### Implementation details

*CROCKETA* is an automated and adaptable *Snakemake* [27] pipeline designed to perform fundamental stages of single-cell analysis for both transcriptomics and repertoire data. The pipeline enables the parallelization of analysis and resources whenever feasible. *Snakemake* facilitates the parallel execution of analysis steps while ensuring that the workflow is carried out sequentially, following a predefined order. The pipeline was developed and tested on Ubuntu 24.04 LTS but is designed to run seamlessly on any standard Linux distribution. For Windows users, it can be easily deployed within a Docker container, ensuring compatibility across different operating systems. We chose *Snakemake* as the workflow management system due to its robust scalability, reproducibility, and ease of containerization. Unlike other workflow managers, *Snakemake* natively supports wrapper integration within Docker, allowing all dependencies to be encapsulated in a controlled environment while maintaining high computational efficiency.

Containerizing the pipeline in Docker offers multiple advantages, including isolation from system dependencies, cross-platform compatibility, and streamlined deployment in cloud or HPC environments. Creating a Dockerized version of the pipeline is straightforward:

1. Define a Dockerfile specifying the required environment (e.g., base image, dependencies, and *Snakemake* installation).
2. Build the image using: `docker build -t pipeline_image`.
3. Run the pipeline within the container using: `docker run --rm -v $(pwd):/workspace pipeline_image snakemake --cores all`

This approach ensures that the workflow remains portable, scalable, and reproducible across diverse computational infrastructures.

The pipeline encompasses multiple stages, including sequencing and cell-specific quality control (QC), alignment, dimensionality reduction, cell clustering, cluster marker extraction, cell annotation, differential expression analysis, functional enrichment, trajectory inference, and RNA velocity. *CROCKETA* incorporates several widely-used single-cell resources, such as *CellRanger* [28], *STAR* [29], *DoubletFinder* [30], *Seurat, SingleR, Azimuth* [31], and *Immunarch*, among others, and stands out in terms of parallelization, flexibility, reproducibility, and customization. The pipeline establishes an environment for each step through *bioconda* and *conda-forge* repositories [32], reducing complexity and enhancing reproducibility.

### Pipeline setup

Workflow configuration relies on three files that allow users to set up the analysis from the outset and specify input files and experimental details. These configuration files include samples.tsv (which contains information about the samples to be analyzed), units.tsv (which provides paths to the input data), and config.yaml (which contains all pipeline-customizable parameters). Input data can be provided in various formats, including FASTQ files and expression matrices.

The pipeline workflow is summarized in the graphical abstract, and each stage of the analysis is detailed below.

### Quality Control

When *FASTQ* files are provided as input and *CROCKETA* is initiated, a quality control (QC) analysis step is executed to asses quality of the sequencing using widely employed tools such as *FastQC* (https://www.bioinformatics.babraham.ac.uk/projects/fastqc/)*[32-a]*. Additional *MultiQC* reports [33] are obtained summarizing *FastQC [34-a]* and *FastQ* screen [34] information and integrating this data with other softwares such as *RSeQC* [35]. When matrix input is employed, this step is skipped.

### Alignment, Data preprocessing & Clustering (Transcriptional primary analysis)

Sequence alignment and quantification are performed only when FASTQ files are provided as input. This step is carried out using *STAR & STARsolo*, ensuring high-performance alignment for single-cell sequencing data. The alignment parameters are fully configurable via the config.yaml file, which serves as the master configuration file for all tools and software within the pipeline, ensuring flexibility and adaptability to diverse experimental setups. While default standard parameters are provided to accommodate common sequencing technologies (e.g., *10x* Genomics, Drop-seq), users can easily modify these settings to tailor the alignment process to their specific dataset. The configuration supports various chemistry versions and multi-omic configurations, including 5’ Chemistry for multi-omic data and 3’ Chemistry for scRNA-seq data exclusively.

Once these steps are performed, all analyses are carried out in *R* environments through *Seurat*.

Regarding primary analysis of transcriptional data, initial preprocessing is required. This preprocessing includes a cell-level QC, in which certain thresholds -such as minimal *UMIs*, mitochondrial expression and others-are user-specified in config file. This step allows for the removal of low-quality or dead cells. In addition, *DoubletFinder* algorithm can be applied to identify and remove hypothetical doublets in each analyzed sample. Retained cells will then undergo normalization, dimensionality reduction (*PCA, UMAP*), clustering & visualization. At the data normalization stage, *CROCKETA* offers flexible handling of batch effects, which is critical given the pipeline’s design to automate and process large-scale datasets. Users can apply standard correction methods or even integrate datasets when batch effect is particularly pronounced.

### Repertoire Data Extraction, Integration & Analysis

If only scRNA-seq data is provided, this step can be skipped if wished.

When input repertoire data is FASTQ formatted, sequencing data are subjected to a second alignment through *CellRanger* pipeline. This enables clonotype data extraction from *scTCR-seq* or *scBCR-seq* technologies. Consequently, a *filtered_contig_annotations*.*csv* output file is generated, containing clonotype data for each cell across all analyzed samples. This file is essential for repertoire integration to single-cell object. In addition, *CellRanger* provides additional QC reports complementary to previously obtained *MultiQC* reports in which we can check *TCR/BCR-*seq quality.

If matrix files are provided as input, contig csv-formatted file must be provided and specified in the corresponding section of the configuration files.

Data integration is accomplished through *scRepertoire R software*. Moreover, based on the specific research focus or the studied disease, users might only be interested in clones not associated with viruses (labeled as non-viral) as happens in a tumoral context. Therefore, clonotype analysis includes a preliminary annotation of TCR clones as viral or non-viral, according to the *VDJ* database (*VDJdb*) [36] through the *Immunarch* R package. Once completed, two different *Seurat* objects are generated: one containing all available information and another containing only non-viral cells, filtering out cells associated with clones matching *VDJdb*. Analysis is conducted on both objects.

At this stage, several repertoire metrics can be obtained in order to elucidate clonal expansion, clonal diversity, clonal overlap and other metrics within each single sample, while also enabling comparisons across all of them.

### Post-clustering Analysis

Finally, post-clustering analyses are applied if enabled by the user to the single-cell object. Analysis starts with a cell annotation step, in which *CROCKETA* aims to elucidate cell identity, and avoid any bias regarding computational software or cell references. With this purpose, two approaches are carried out:

- <u>Automatic annotation:</u> Up to three softwares of analysis can be employed: *SingleR, Azimuth & scType* [37]. *SingleR & Azimuth* employ pre-defined references (more than one reference can be specified at once), while *scType* employ a user-defined reference provided in their documentation which is very easy to edit to, for example, add missing cell populations, tissues or gene markers.
- <u>Manual data explore:</u> A gene-set of interest is user-provided and expression for every gene is assessed in our single-cell object through *Seurat* means. This information is complementary to automatic annotation approach to confirm annotation results. If there is no gene-set specified, a general list of marker genes regarding bone-marrow subpopulations is provided.

Results are extracted at both single cell level and cluster level. The main point is to integrate all cell-label information in order to provide a robust cell annotation.

Furthermore, this post-clustering analysis will enable population marker extraction through differential expression analysis, up/down-regulated biological pathways from functional enrichment analysis, and different cell states in a pseudotime space from trajectory inference analysis. These analyses are performed using state-of-the-art softwares including *Vision, GSEApreranked* method from *msigDB* [38], *Slingshot* and *Velocyto* [39].

**[Graphical abstract]:** Schematic representation of *CROCKETA* workflow.

## RESULTS

Several output files are generated throughout the *CROCKETA* workflow.

The pipeline outputs aggregate quality control reports from multiple stages of the processing in a *multiQC* report. Regarding single-cell analysis, a wide variety of pdf-formatted visualizations are generated for each stage including UMAP visualizations, Seurat-based figures, clonality-descriptive plots and others. Additionally, each analysis stage performed in *Seurat* provides its respective single-cell object in .rds format for subsequent processing. Moreover, excel-formatted files are also created to provide statistical results. Primary stage outputs are depicted at online documentation: https://github.com/OncologyHNJ/crocketa. Some further transcriptomic and clonal results are described below.

### Repertoire Analysis

Once transcriptomics data is properly processed up to the cell clustering step (Fig. 1A), clonal data is integrated and analyzed. Initially, clonal single-cell data is summarized in UMAP coordinates, where clonal cells are ranked according to certain user-defined clonal frequency ranges (relative or absolute values): *Hyperexpanded, Large, Medium, Small* and *Single*. Bar plots illustrating clonal diversity results, cell counts, relative proportions and unique number of clones, are generated (*Fig. 1B)*. In the case of TCRs, viral annotation is performed as detailed in the methods section, utilizing *VdJdb*, and several visualizations are generated for alpha-chain, beta-chain and paired-chain viral annotated TCRs (*Fig. 1C*).

**Figure 1.**
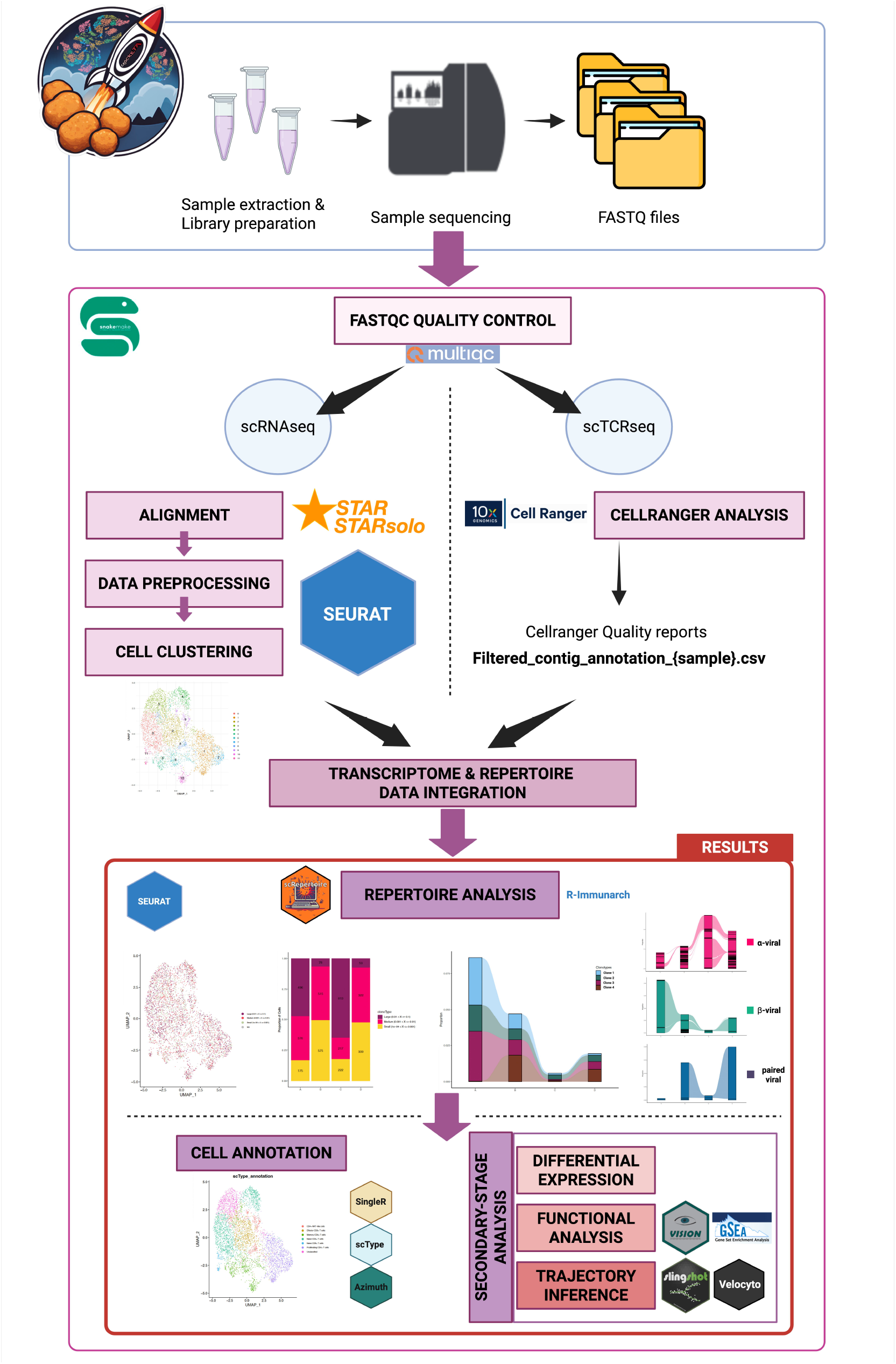
Immune Repertoire analysis. A) *UMAP* dimensionality plots colored by condition of interest [left] and cell cluster [right]. B) Immune repertoire dataset. *UMAP* dimensionality plot colored by clonal frequency [left]; Bar plot showing the proportion and number of cells per clonal frequency range in each condition of interest [right]. C) TCR viral-annotated clones. Fraction of viral annotated clones per condition per chain: α or β [left] and alluvial visualization of viral clones per chain and per condition [right] D) Most frequent clones per condition through bar plots (Top 100) and *UMAP* dimensionality plots (Top 5). E) Clonal overlapping across conditions of interest through Upset-like plot and *UMAP* dimensionality plot. F) Frequency fluctuation of common clones across conditions of interest through alluvial plot.

Following TCR viral annotation, the same analyses are applied to both the full single-cell object and the non-viral single-cell object. Statistical graphics and tables are extracted.

Some per-sample results can be generated to elucidate the most frequent clones, visualized in either bar plots or UMAP visualizations (*Fig. 1D)*. Additionally, results concerning clonal overlap among samples or conditions of interest are reported. Venn diagrams and/or upset-like plots are provided for one-to-one and all-to-all comparisons (*Fig. 1E)* in order to observe all possible combinations across conditions simultaneously through *ggvenn* (https://cran.r-project.org/web/packages/ggvenn/readme/README.html) and *UpsetR* software [40]. Further visualizations are included to identify common clones to all conditions (e.g. UMAP plots, *Fig. 1E, 1F*), in addition to an alluvial plot illustrating frequency fluctuations across conditions for each individual sample/condition *(Fig. 1F)*. More results and updates will be provided on the online documentation.

### Post-clustering Analysis

Upon completion of the repertoire analysis, *CROCKETA* proceeds to the final stage of the workflow, which, while not essential, serves as an optional secondary step in the transcriptional analysis. Output for differential expression analysis, functional enrichment, and trajectory inference may be computed. *CROCKETA* includes an intermediate cell annotation step in order to elucidate cell identity. Several visualizations will be provided for each of the genes of interest specified through Seurat-based methods such as *FeaturePlot* graphics (*Fig. 2A*). Automatic-annotation results are extracted from every software and reference applied at a cell or cluster level, and *UMAP* plots might be extracted for each approach (*Fig. 2B*). Through the combination of all possible annotation results from the specified approaches, user can finally assign a robust label to each of the cell/clusters in the assay.

**Figure 2.**
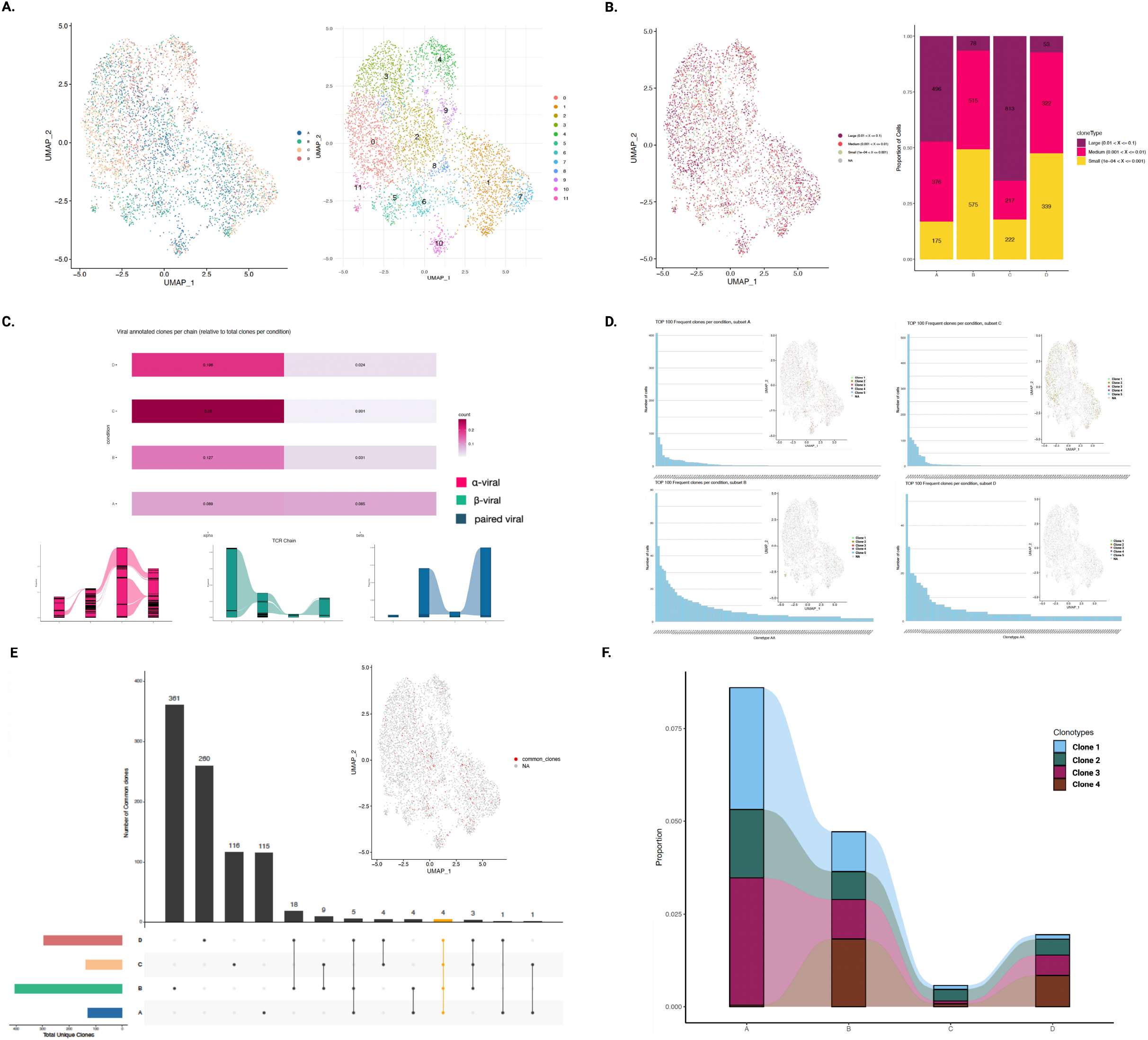
Cell annotation. A) Expression of *MKI67* gene, marker of proliferation. B) *scType* preliminary annotation results. C) *Vision* results for Hallmark_E2F_targets. D) *Velocyto* results to elucidate individual cell future states. Colored according to cell cluster. E) *Slingshot* trajectory inference results. Colored according to cell cluster. F) *Slingshot* trajectory inference results. Genes influencing trajectory 2: from cluster 7 (yellow) to cluster 3 (green).

To explore functional enrichment, several methods might be applied to highlight transcriptional activation of biological pathways. According to *Vision*, the expression of *E2F*-targets (based on *Hallmarks*) seems to be enriched in clusters 1 and 7 (*Fig. 2C*), consistent with MKI67 expression as a proliferation marker and with preliminary *scType* annotation results, identifying these cells as proliferating CD4+ T cells. Regarding trajectory inference results, several visualizations can be extracted to elucidate cell-state transitions through different approaches *(Fig. 2D, 2E, 2F)*.

### Example use-case

The *CROCKETA* pipeline has been tested with real patient samples from pediatric oncology cases, including sarcomas and neuroblastomas. In these cases, tumor-infiltrating lymphocytes (TILs) were isolated and co-cultured. scRNA-seq along with TCR sequencing was carried out to characterize the immune repertoire at a high resolution. The data generated from these analyses are currently being prepared for publication, and the results hold significant scientific and clinical value, particularly in understanding tumor-immune interactions and identifying potential therapeutic targets. Here we provide partial results for a subset of the sequenced cells. Results for this subset are extracted and visualized in Figures 1 and 2.

This example dataset does not include BCR data, but every step of analysis can be applied to BCR and TCR datasets independently or jointly. The only step not implemented yet for BCRs is the viral clone annotation step, which is currently available only for T-cells.

All procedures involving human participants were approved by Ethic Committee under protocol number R-0076/19, and written informed consent was obtained in accordance with institutional guidelines and regulations.

### Strengths and Limitations

*CROCKETA* integrates a carefully selected set of widely recognized, high-impact tools (e.g., *Seurat, Immunarch, STAR*) that have been benchmarked in peer-reviewed studies, ensuring both methodological rigor and reproducibility. The pipeline is based on well-established and validated approaches, combining flexibility and modularity to accommodate a wide range of research contexts, from fundamental immunology to translational oncology. Researchers can analyze transcriptomic and repertoire data separately or together, with customizable workflows designed to prioritize specific research objectives. *Snakemake’s* parallelization further enhances the pipeline’s scalability, making it suitable for large-scale cohorts. By automating the integration of clonal and transcriptomic data, *CROCKETA* significantly reduces analytical variability and accelerates hypothesis generation. The pipeline’s automated multi-step integration (e.g., clonotype-transcriptome mapping) minimizes manual biases, enabling faster and more reliable hypothesis testing. This approach is valuable across a broad spectrum of applications, from basic immunology studies to the discovery of clinical biomarkers in oncology and immunotherapy research.

However, *CROCKETA* has some constraints. Its reliance on CellRanger for BCR/TCR alignment limits repertoire analysis to *10x* Genomics datasets, excluding non-*10x* platforms like SMART-seq, though scRNA-seq analysis remains feasible. Additionally, while *Snakemake’s* resource management aids in handling large datasets, the pipeline still inherits the computational demands of single-cell tools, such as Seurat’s memory-intensive clustering, which requires high-performance computing (HPC) or cloud environments for large-scale analyses (>1,000,000 cells). Furthermore, the integration of clonotype and transcriptome data assumes high-quality input, meaning that low RNA-seq library complexity or sparse TCR/BCR reads can lead to incomplete or inaccurate cell-clonotype pairings.

## DISCUSSION AND CONCLUSION

*CROCKETA* represents a versatile, robust and scalable solution for the integrated analysis of transcriptional and clonal data at the single-cell level. Built on *Snakemake*, it ensures parallelization, reproducibility, and modular customization, addressing key challenges in multi-omic single-cell studies. Unlike fragmented approaches, *CROCKETA* streamlines the entire workflow, from raw data processing to advanced immune repertoire characterization, facilitating the identification of expanded T- and B-cell clones and their transcriptional states. Its seamless integration of state-of-the-art tools for clonal annotation, transcriptomic signatures, and visualization makes it a powerful resource for researchers exploring immune dynamics in cancer, autoimmunity, and infectious diseases.

Designed with flexibility and scalability in mind, *CROCKETA* is compatible with diverse experimental designs and sequencing technologies. The pipeline will undergo continuous updates, incorporating new analytical features and improved compatibility with emerging methodologies. By enabling the correlation of immune clonality with functional states, *CROCKETA* has the potential to contribute significantly to immunotherapy research, particularly in identifying clonal expansions linked to exhaustion or memory phenotypes.

## Availability of data and materials

*CROCKETA* pipeline, documentation and source code are available in the Github repository: https://github.com/OncologyHNJ/crocketa

## Key Points

- *CROCKETA* is the first *Snakemake*-based pipeline to automate the integration of single-cell transcriptomics with BCR/TCR repertoire data, enabling simultaneous exploration of clonal expansion and transcriptional heterogeneity within a unified workflow.
- Leveraging *Snakemake’s* parallelization and *Conda* environments, *CROCKETA* ensures reproducible, scalable analysis from raw FASTQ files to publication-ready visualizations, adaptable to studies ranging from small cohorts to large-scale datasets.
- The pipeline provides granular clonal metrics (diversity, overlap, expansion) and unique features like automated viral TCR annotation via *VDJdb*, bridging clonality with antigen-specific immune responses.
- Users can tailor analyses through customizable configuration files, selecting specific tools (e.g., *Azimuth, SingleR*) and adjusting parameters for QC, clustering, or trajectory inference without modifying core code.
- By correlating clonal dynamics with transcriptional states (e.g., exhausted T-cells in cancer),

*CROCKETA* unlocks actionable insights into disease mechanisms, vaccine responses, and immunotherapy efficacy.

## ACKNOWLEDGEMENTS

The authors sincerely appreciate the work of the whole FIBHNJS staff.

---

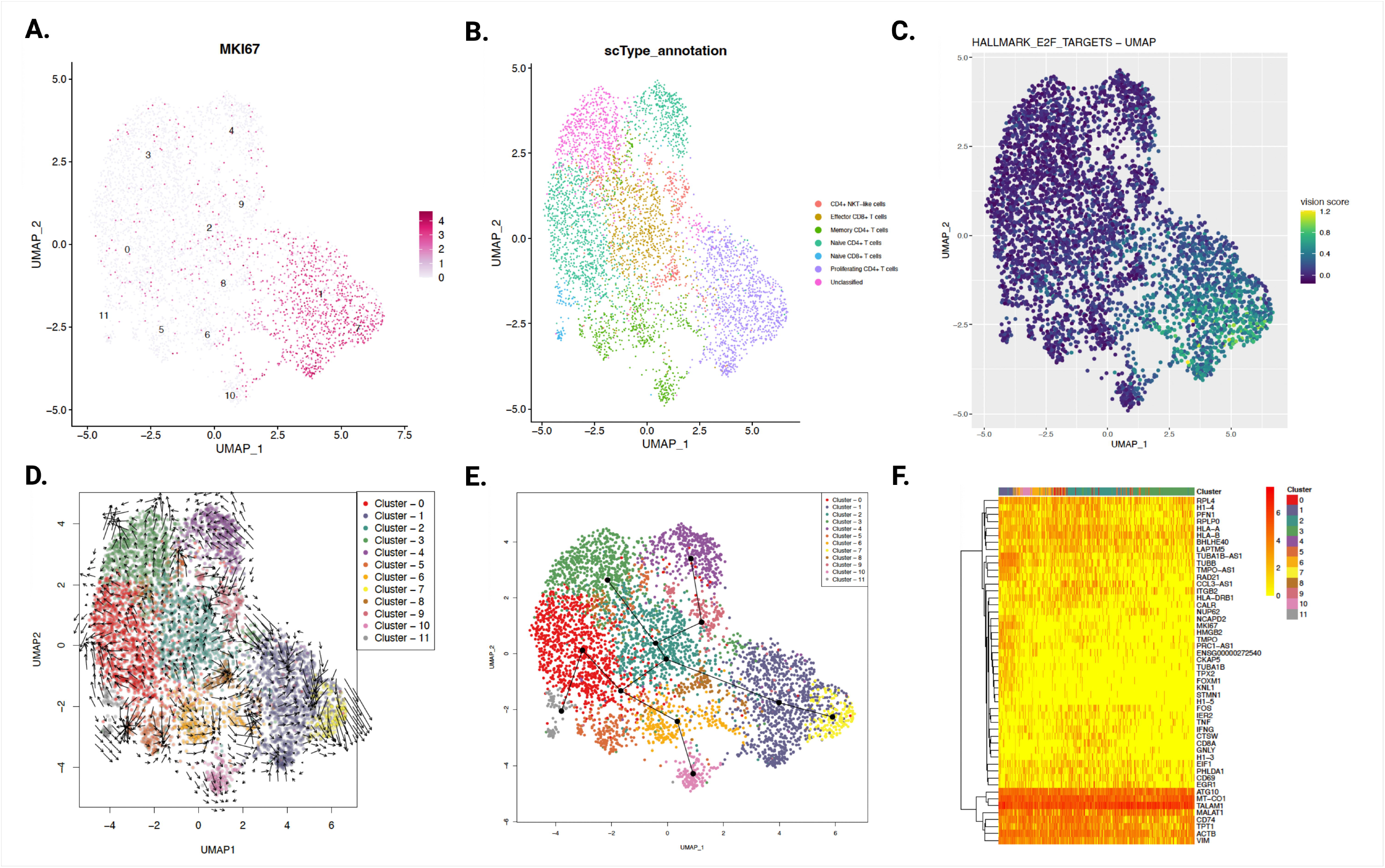

